# Src42A is required for E-cadherin dynamics at cell junctions during *Drosophila* axis elongation

**DOI:** 10.1101/2022.04.20.488907

**Authors:** L. Chandran, W. Backer, H.B. Beati, D. Kong, S. Luschnig, H.-Arno J. Müller

## Abstract

Src kinases are important regulators of cell adhesion. Here we explored the function of Src42A in junction remodelling during *Drosophila* gastrulation. Src42A is required for tyrosine phosphorylation at bicellular and tricellular junctions in germband cells and localizes to hotspots of mechanical tension. The role of *Src42A* was investigated using maternal RNAi and by CRISPR-Cas9-induced germline mosaics. We find that during cell intercalations, *Src42A* is required for the contraction of junctions at anterior-posterior cell interfaces. The planar polarity of E-cadherin is compromised and E-cadherin accumulates at tricellular junctions after *Src42A* knock down. Furthermore, we provide evidence that Src42A acts in parallel to Abl, which has also been implicated in cell intercalations. Our data suggest that Src42A is involved in two related processes, one affecting tension generated by the planar polarity of MyoII and secondly, it can act as a signalling factor at tAJs in controlling E-cadherin residence time.

## Introduction

Gastrulation represents a crucial morphogenetic process in early embryogenesis by which the blastomeres enter cell fates in three different germ layers and the principal body axes become established (Williams and Solnica-Krezel 2017). Much of the cellular basis of gastrulation movements is well known from studies in model organisms, including *Drosophila melanogaster* (Gheisari et al. 2020; Paré and Zallen 2020). In *Drosophila* the anterior-posterior (AP) pattern originates during oogenesis to set up molecular gradients in the fertilized egg, which inform the elongation of the AP axis in a process called germband extension (Kong et al. 2017). The translation of the AP patterning into directional movement of the germband critically involves the generation of planar cell polarity through Toll-like receptor proteins (Paré et al. 2014).

A band of ventral-lateral epidermal cells forms the germband, which during gastrulation more than doubles its length along the anterior-posterior direction of the embryo, while narrowing in the dorsal-ventral direction (Kong et al. 2017). Three different modes of cell behaviours driving cell intercalation during the extension of the germband have been described: T1-transition, multiple rosette formation and sliding vertex (Bertet et al. 2004; Blankenship et al. 2006; Vanderleest et al. 2018). All these three behaviours require adherens junctions (AJs) to undergo precisely controlled remodelling processes (Rauzi et al. 2010; Levayer et al. 2011; Levayer and Lecuit 2013). According to their position within the blastoderm epithelium, AJs can be classified into bicellular junctions (bAJs) describing the 2-cell interface and tricellular junctions (tAJs) describing a cell vertex where three cells attach to each other; both, bAJs and tAJs are modulated during germband elongation.

The T1 transition describes the intercalation between four cells that exchange their neighbours by shrinking the bAJs in vertical direction and extending a new junction in horizontal direction (Bertet et al. 2004). The localized contraction and extension of the bAJ is facilitated through a planar polarized distribution of non-muscle myosin II (MyoII), whose levels are enriched at vertical (AP) cell interfaces, while the scaffolding protein Bazooka (Baz) along with the E-Cadherin-Catenin complex is enriched at the horizontal (dorsoventral, DV) interfaces (Paré and Zallen 2020; Kong et al. 2017; Zallen and Wieschaus 2004). Interference with the planar polarization of either of these proteins negatively affects cell intercalation. For example disruption of the AP patterning via *Krüppel* RNAi knockdown affects junctional remodelling and MyoII AP planar polarity (Bertet et al. 2004). Junctional remodelling is also dependent on tyrosine phosphorylation; for instance, the DV planar polarity of β-catenin is compromised after Abelson (Abl) kinase knockdown (*abl^i^*), resulting in a delay of cell intercalations (Tamada et al. 2012).

More recent studies have shown that tAJs also play a role in germband extension. The transmembrane protein Sidekick (Sdk) exclusively localizes at the tAJs and is vital for maintaining the length of the AP and DV interfaces. *Sdk* mutants show defects in apical vertex adhesion and less strain in the tissue which affects the rate of T1-transition significantly (Finegan et al. 2019). In addition, Sdk localization changes when there is increased mechanical tension. Expressing Myosin Light chain phosphatase resulted in less Sidekick accumulation at the tAJs suggesting that of Sidekick protein levels are controlled by mechanical tension (Letizia et al. 2019). Recently, it was shown that Canoe and its vertex localization is vital for cell intercalation; the mobility of Canoe from the tAJs to bAJs is inevitable for germband extension and this mobilization is mediated by phosphotyrosine signalling through Abl kinase (Yu and Zallen 2020).

In addition to Abl, Src Kinases have also been shown to be required for germband elongation. A first report provided evidence that Src42A, in conjunction with the small GTPase Rac1, is required for the formation of actin-rich protrusions on the basal side of multicellular rosettes (Sun et al. 2017). More recently it was shown that Src42A is a component of a signalling pathway mediating the cell surface information from Toll-2 receptor towards PI3-kinase in order to promote the formation of MyoII planar polarity (Tamada et al. 2021). However, despite the well-known functions of Src family kinases in cell adhesion, the role of Src42A in regulating the distribution and dynamics of the E-cadherin adhesion complex has not yet been studied.

In this work we explored the function of one of the two Src kinases encoded in the *Drosophila* genome, *Src42A*, in fine tuning both tAJs and bAJs during T1-transition. Consistent with previous reports, we find that Src42A has a substantial impact on T1 transitions. After Src42A single knockdown (*Src42A^i^*) the contraction of the AP border and the planar polarity of the bAJs are impaired. The delayed T1-transitions in *Src42A^i^* embryos slowed down cell intercalation and correlated with reduced tension at the AP cell border and a delay in the rate at which the AP borders shrink during T1 transition. *Src42A^i^* embryos exhibit increased E-cadherin level at the AP cell border suggesting that the turnover of E-cadherin is affected.

Consequently, E-cadherin intensity at the tAJs is increased in intercalating cells. E-cadherin dynamics at tAJs show an accumulation and dispersion cycle, which is affected by reducing the level of Src42A. Our data suggest that Src42A is involved in two not mutually exclusive processes; in addition to its function in Toll-2 like receptor signalling instructing the planar polarity of MyoII, we propose that it also acts as a signalling factor in controlling E-cadherin residence time at tAJs.

## Results

### Dynamic association of Src42A with plasma membrane domains in the early embryo

The *Drosophila* genome encodes two Src homologs, Src42A and Src64B. To detect endogenous Src42A protein, we generated an antiserum against a full-length GST-Src42A fusion protein. Src42A levels were significantly reduced in *Src42A^26-1^* homozygous embryos compared to controls (Fig. 1A,B; (Fig. S1A to E)). Immunostaining of embryos expressing HA-tagged Src42A or Src64B revealed that the antibody was specific for *Src42A*, while no cross-reactivity with Src64B was detected (Fig. S2). Consistent with earlier studies we found Src42A to be associated with the plasma membrane in the early embryo (Takahashi et al. 2005). Src42A exhibits a differential accumulation at distinct plasma membrane domains during cellularization. In early to mid-cellularization, Src42A was localized at the apical aspect of the ingressing plasma membrane, where it partially colocalized with Baz at the plasma membrane cortex (Fig. 1C,C’). A second accumulation of Src42A was present at the basal tip of the ingressing furrow canal, where the localization overlapped with Myo II (Fig. 1C,C’). During late cellularization, when spot AJs move apically, Src42A accumulated at the emerging apical AJs, whereas levels of Src42A at the furrow canals decreased (Fig. 1C,C’). During gastrulation, Src42A was observed at bAJs and tAJs on the apical side of the lateral epidermis during germ-band extension. (Fig. 1D,D’). Src42A was slightly enriched at the AP interface where Myo II was localized during the generation of planar cell polarity (Fig. 1D’ stage 7 embryos). We conclude that by using a specific antibody, we detected a differential localization of Src42A at distinct plasma membrane domains in early embryos. The observation that Src42A was especially enriched at tension-generating hotspots in between the cells prompted us to examine the function of Src42A during gastrulation.

**Figure 1:**
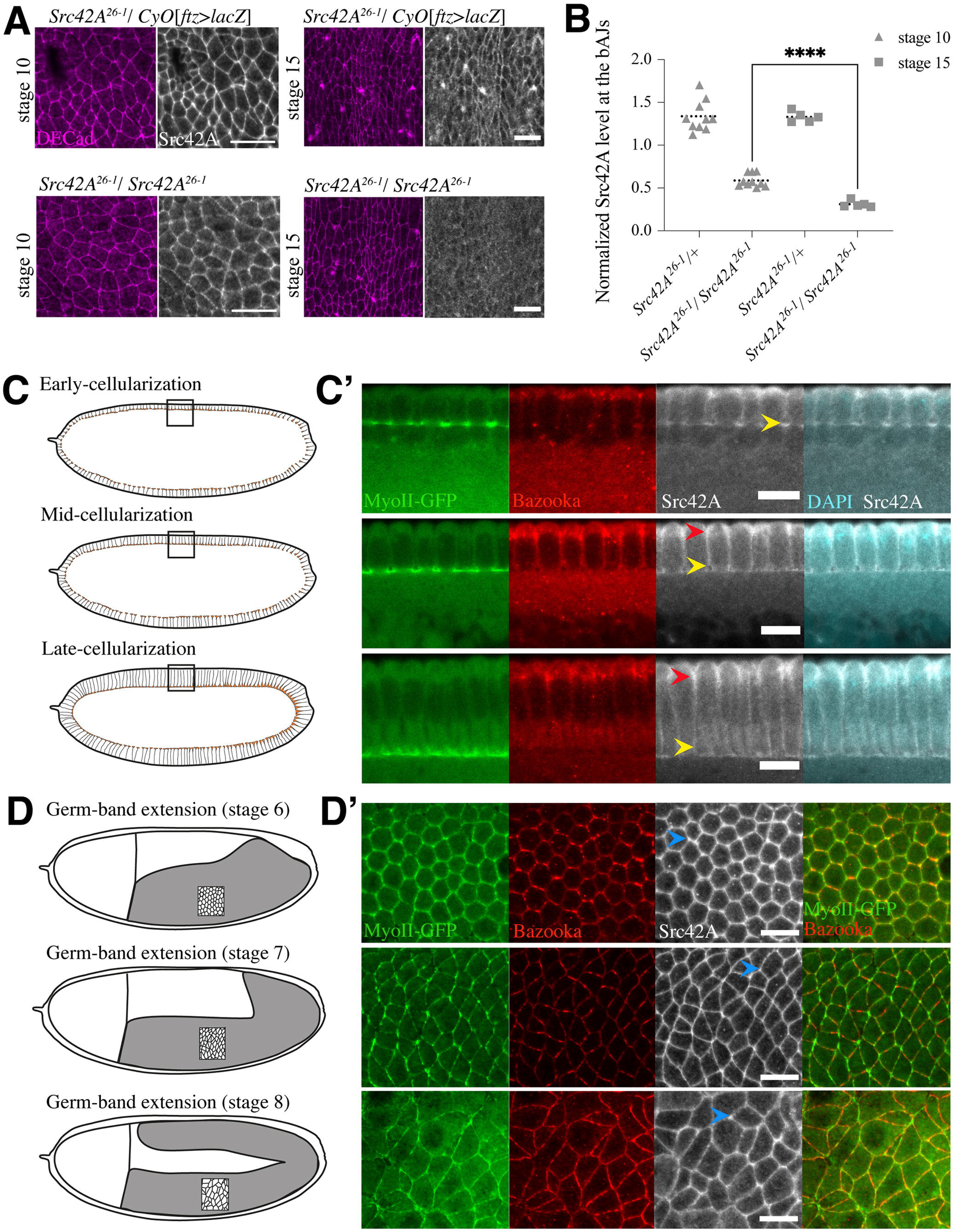
Subcellular localization of Src42A during cellularization and germband extension. **(A)** Confocal imaging of heterozygous *Src42A^26-1^*/*CyO*[*ftz>lacZ*] and homozygous *Src42A^26-1^*/ *Src42A^26-1^* embryos stained for E-cadherin (magenta) and Src42A (grey), embryonic stages are mentioned on the left side of the image sets. **(B)** Maternal Src42A quantification at bicellular adherence junctions in stage 10 and stage 15 embryos, plotted value showing Src42A intensities were normalized over E-cadherin intensity. Each data point (a grey triangle or square) in the graph corresponds to average pixel intensity measured in 10 cell junctions in a single embryo. Dotted line represents mean value and error bars are drawn using range. **(C,D)** Schematics of cellularization and germband extension stages corresponding to the stages in **(C’,D’). (C’,D’)** Confocal micrographs of *MyoII::KI-GFP* (Ambrosini et al. 2019)embryos in cellularization (in transversal section) and germband extension (surface projection) stages. Embryos were stained for Bazooka (red), Src42A (grey), DAPI (cyan) and MyoII-GFP (green). Scale bars corresponds to 10μm. Arrow heads marking furrow canal (yellow **C’**), adherens junction (red **C’**) and cell vertex (blue **D’**).

### Maternal knockdown of *Src42A* affects embryogenesis from the blastoderm stage

Early embryos homozygous mutant for Src42A harbour considerable amounts of maternal Src42A protein (Fig. S1E). The elimination of maternally supplied Src42A is not possible by the conventional autosomal DFS-FRT technique, since the Src42A gene locus is present between the centromere and the most proximal available FRT site (Chou and Perrimon 1996). Therefore, to deplete maternal Src42A, RNAi knock-down experiments were conducted by employing the UAS/GAL4 system driving a transgenic short hairpin RNA using the *P{TRiP.HMC04138}* line (Staller et al. 2013; Brand and Perrimon 1993) (Fig. 2A). Embryos expressing *TRiP04138* showed reduced hatching rates with variable penetrance and expressivity depending on the maternal Gal4 driver used (Fig. 2B). A combination of *P{mat 4-GAL4-VP16}67; P{mat 4-GAL4-VP16}15* (called *mat67; mat15* hereafter) was most efficient as embryos showed about 80% reduction in hatching rates and were hence used for all the experiments (called *Src42A^i^* in this study). *Src42A^i^* embryos displayed variable cuticle phenotypes, which were stronger compared to zygotic *Src42A^26-1^* homozygous mutants (Fig. 2E). In cleavage stages, 40% of *Src42A^i^* embryos exhibited an irregular interface between the central yolk and the clear cortical cytoplasm (Fig. 2B, D). By immunoblotting, we found that embryos with defects in the cortical cytoplasm exhibited the most severe reduction in Src42A protein levels (Fig. 2C). Similar defects were also observed in embryos injected with dsRNA against Src42A mRNA (Sun et al. 2017). Since the Src42A knockdown was more pronounced in embryos with defective cytoplasm, these embryos were used for further analysis.

**Figure 2:**
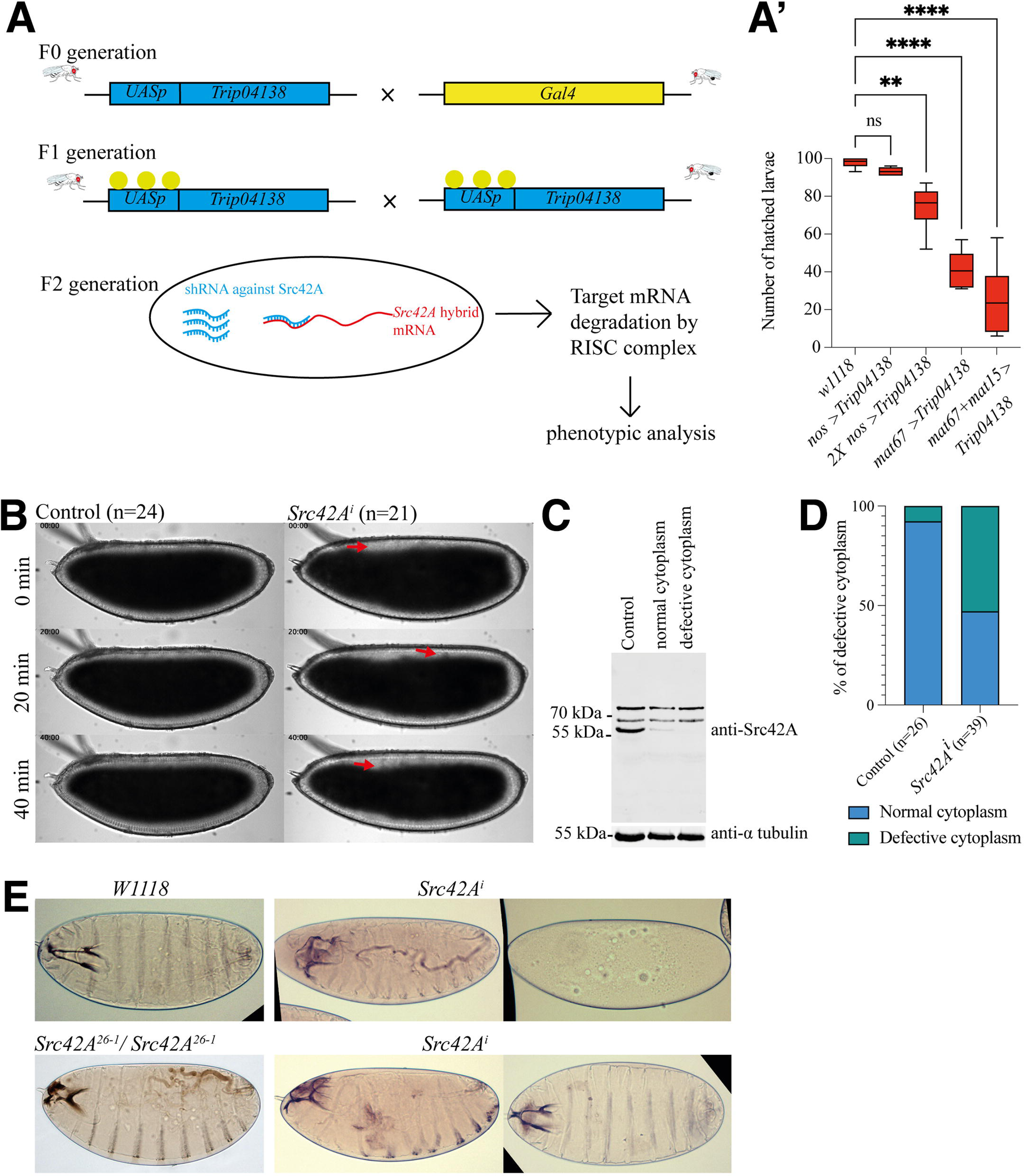
Characterization of Src42A knockdown. **(A)** A schematic view shows crossing scheme for Src42A knockdown using UAS/GAL4 system. *Trip04138* expresses short hairpin RNA (blue) against endogenous Src42A (the target mRNA in red). *Gal4* shown in yellow. (**A’)** Src42A knockdown analysis using different maternal *Gal4* driver lines, statistics were done using Dunnett’s multiple comparison test shows significant difference among the mean values. The p values of *nos*>*Trip04138*, 2X*nos*>*Trip04138*, *mat67*>*Trip04138* and *mat67*+*mat15*> *Trip04138* are 0.8788(ns), 0.0043(**), 0.0001(****) and 0.0001(****) respectively. **(B)** Bright field images showing defective cytoplasm during cellularization stage in control and *Src42A^i^*. Red arrows indicate defective cytoplasm. (**C)** Western Blot analysis to genotype *Src42A^i^*. Embryos (15 embryos each) showing defective and normal cytoplasm from *Src42A^i^* crosses were selected, lysed under denaturing conditions, and subjected to SDS-PAGE and Western blot analysis. Embryos with defective cytoplasm show strong reduction in Src42A level (∼59kDa). As a loading control alpha tubulin(∼55kDa) is used. (**D)** Percentage of defective cytoplasm were calculated based on the total n values (number of embryos counted). **(E)** Cuticle preparation from wild type (*w^1118^*) and *Src42A* zygotic mutant (*Src42A^26-1^*/*Src42A^26-1^*). *Src42A^i^* cuticles showing stronger phenotypes.

### Src42A is required for germband extension and for normal phosphotyrosine levels at bAJs and tAJs

Our immunostaining experiments revealed that during germband extension Src42A was localized at bAJs and tAJs (Fig. 1B). bAJ- and tAJ-resident proteins have been reported to undergo dynamic redistributions during germband extension (Uechi and Kuranaga 2019; Finegan et al. 2019). *Src42Ai* embryos exhibit defects in germband elongation, when analysed by video time-lapse recordings (Fig. 3A). Plotting of cumulative displacement length of the germband over time revealed a substantial delay in its AP extension (Fig. 3B). Germband cells showed a 0.25- and 0.16-fold delay in *Src42A^i^* embryos for the fast and the slow phase, respectively (Fig. 3C,D,E). We further examined germband cells in *Src42A^i^* embryos at a subcellular level. Src42A has several known phosphorylation targets, which are present in bAJs and tAJs. One of the best known targets is β-catenin encoded by the *armadillo* (*arm*) gene in *Drosophila*, which is mainly localized at bAJs (Takahashi et al. 2005; Brunet et al. 2013). Indeed, at the onset of germband extension, (stage 7), *Src42A^i^* embryos exhibited significantly reduced phosphotyrosine levels at bAJs and tAJs (Fig. 3F). This indicates that Src42A is required for normal levels of protein phosphorylation at tyrosine residues at bAJs and tAJs. Consistent with earlier findings that Src42A controls germ band elongation by changing the properties of signalling factors at the subcellular level (Sun et al. 2017; Tamada et al. 2021), these results demonstrate that Src42A-dependent tyrosine phosphorylation is required for normal germband elongation.

**Figure 3:**
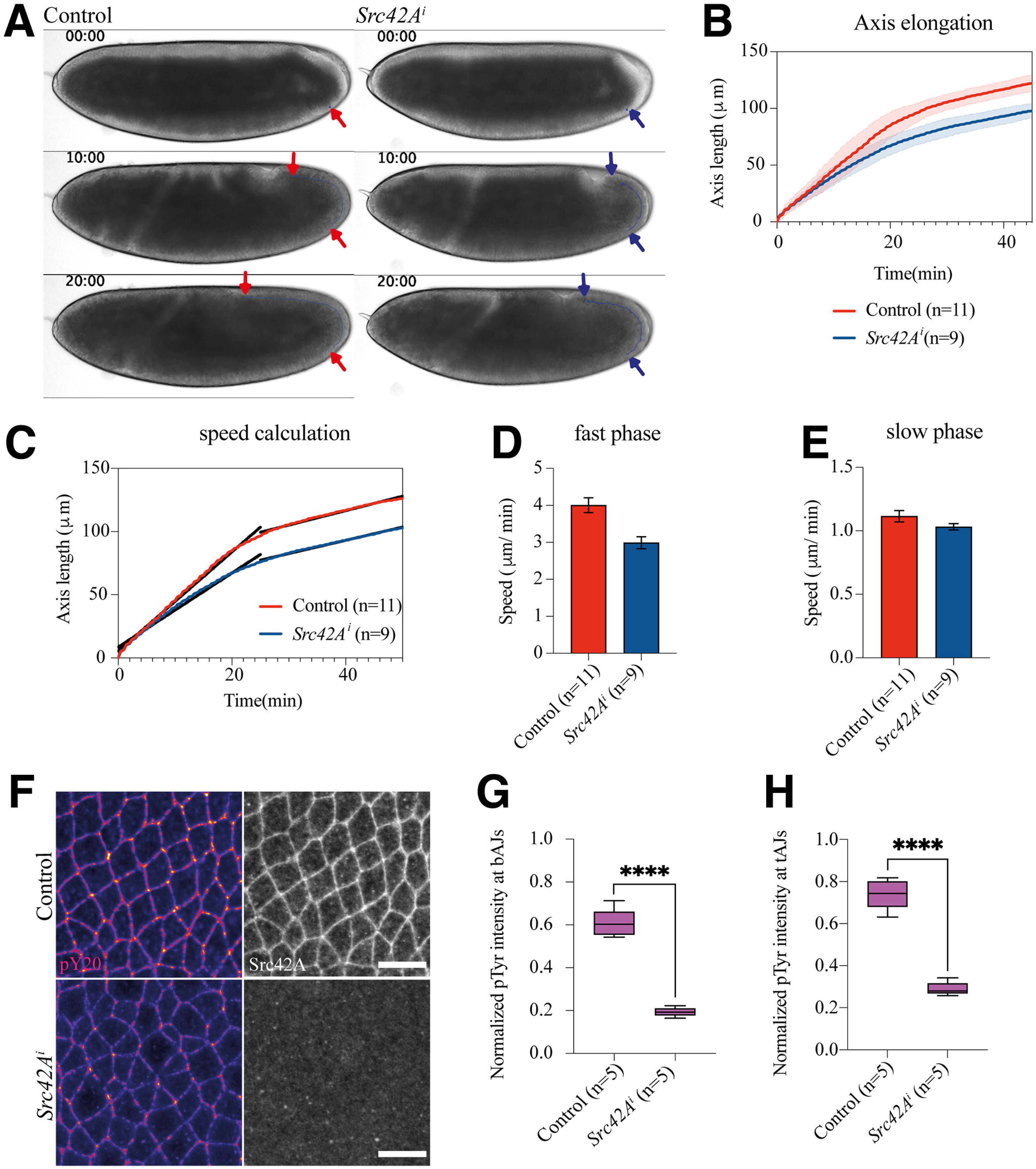
Knock-down of Src42A affects the speed of germband extension. **(A)** Bright field still images at different time points obtained from time-lapse movies of control and *Src42A^i^* embryos during germband extension. Arrow heads are marking the length of elongation at different time points (time indicated in mm:ss format). **(B)** Cumulative length of the extending germband was calculated using the displacement values and plotted over time. **(C)** Linear regression analysis was performed for the axis elongation data for fast and slow phase of germband extension and slope values of the linear curve (black) was considered as the speed of the germband. **(D,E)** Average speed values from the embryos were plotted for fast and slow phases of the germband elongation. The error values are inferential based on confidence intervals (CI). The n value represents the numbers of embryos and R^2^ is <0.5 showing the goodness of fit. **(F)** Germ-band cells undergoing intercalation on the subcellular level (the scale bar indicates 10µm). Maximum intensity projection (0.8µm deeper from the surface) of phosphotyrosine antibody (pY20) stained embryos along with Src42A staining. **(G,H)** Control and *Src42A^i^* show significant difference in the pY20 signal at bicellular and tricellular junctions (bAJs & tAJs), p value is <0.0001.

### Src42A is required for normal timing of T1 transitions

T1 transitions are a key cell behaviour during the fast phase of germband extension. It involves junction contraction at the AP cell interfaces followed by neighbour exchange along the DV axis resulting in tissue elongation (Bertet et al. 2004). Recently, it was shown that Src kinases play an instructive role in planar polarizing MyoII and Baz downstream of Toll-2 (Tamada et al. 2021); when both Src42A and Src64B were knocked down, the rate of intercalation determined by altered arbitrary cell edge contraction rates was reduced. Based on the finding that *Src42A^i^* embryos display drastic reduction in the fast phase of germband elongation (Fig. 3D), we sought to investigate T1 transitions in *Src42A^i^* embryos. Using Utrophin-GFP as a marker for the plasma membrane associated actin-cortex, T1 transitions were recorded in control and *Src42A^i^* embryos. In 4-cell vertices undergoing T1-transitions, the shrinking junction along the AP axis strongly reduces its length with an average of 1.25 minute (Fig. 4A). When compromising *Src42A* function, the T1 process was delayed (Fig. 4B). In the large majority of T1 transitions in *Src42A^i^* embryos, the AP junction was not markedly reduced after 1 minute (Fig. 4C,D). This indicates that the rate at which T1-transitions occur is delayed due to less contractility at the AP junctions. We conclude that *Src42A* is required for contraction of AP junctions during T1 transitions in germband elongation.

**Figure 4:**
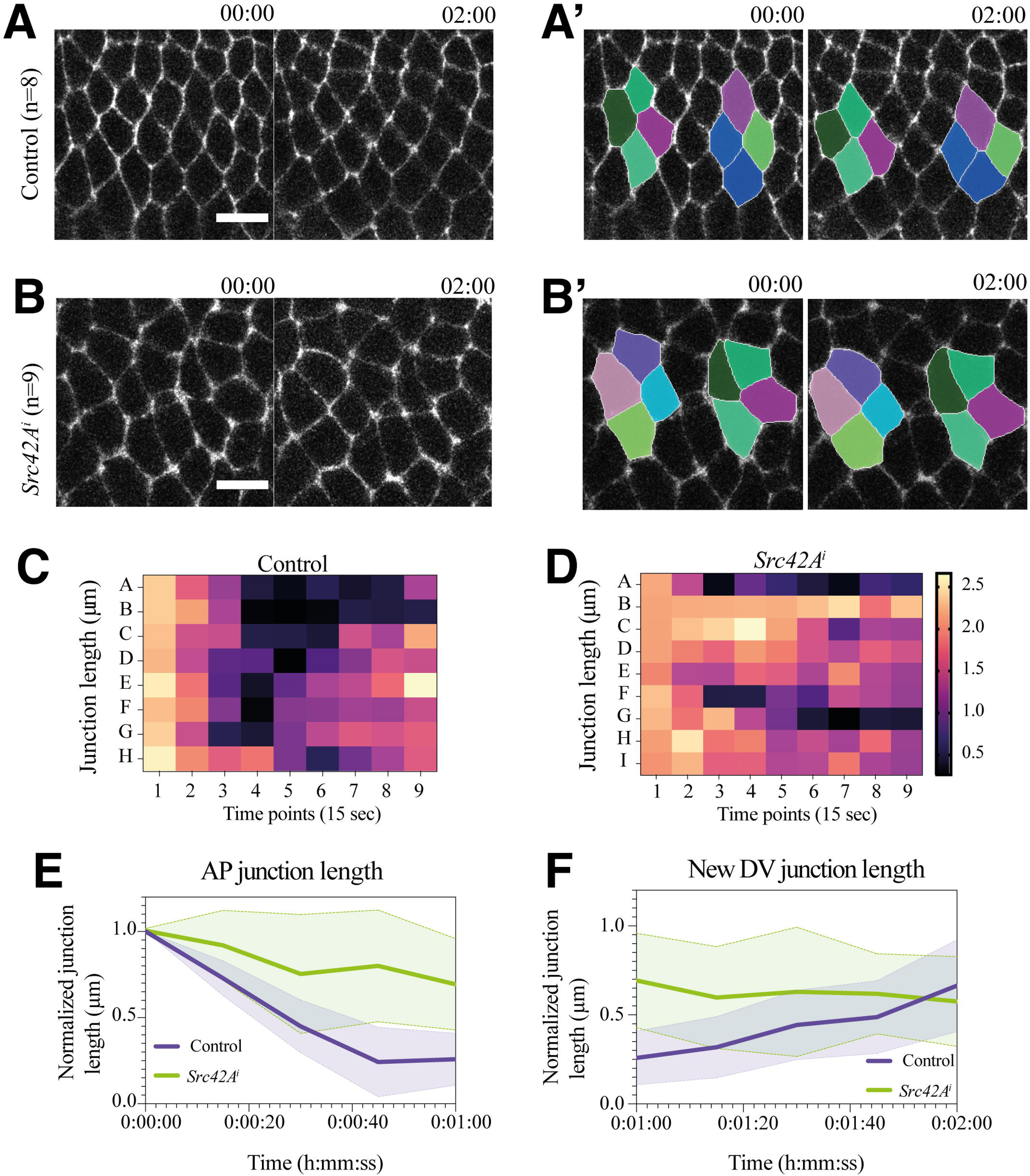
Delay of T1 transitions in germband extension of *Src42A^i^* embryos. **(A,B)** Control and *Src42A^i^* embryos expressing Utrophin-GFP: T1 transitions are indicated (timepoints are depicted on top of the images (in mm:ss). (**A’,B’)** Segmented images, single cell group undergoing T1 transition (marked in colours). **(C,D)** Heat map shows raw data (junction length), x-axis shows nine time points at 15 seconds intervals. Y-axis shows individual cell groups (A, B, C etc.,) undergoing T1 transition. **(E)** Normalized AP border length plotted over time. **(F)** DV border length is plotted over time. Both shrinking and new junction length were normalized to initial length at first time point consisting of 2-2.5µm length. (*n* equals the number of T1 cell groups counted for analysis among 5 embryos). At least one cell group is considered for the analysis of single embryo. Unpaired t-test with Welsch correction shows p value of 0.0357 (less than<0.05) indicating the significant difference between control and *Src42A^i^*.

In T1 transitions, the AP cell border is enriched with MyoII, which mediates tension through its contractile properties. Moreover, a supracellular enrichment of MyoII was also reported to occur due to tensile forces along the AP cell interfaces (Fernandez-Gonzalez et al. 2009; Bertet et al. 2004). In *Src42A*, *Src64B* double knockdown embryos, MyoII planar polarity was severely reduced (Tamada et al. 2021). We found that in single knockdown embryos for Src42A (*Src42A^i^*) T1 transitions are delayed and we asked whether this correlates with a reduction in MyoII-dependent tension at the AP border of the cells undergoing T1 transition. To measure the tension at the AP cell boundaries, we performed laser ablation experiments in intercalating cells and measured the recoil velocity. When the AP border was cut, the detached tAJs moved slower in *Src42A^i^* embryos compared to control (Fig. 5A). The displacement of tAJs were normalized with initial length and plotted over time (Fig. 5C). A non-linear regression analysis on displacement data revealed that the initial recoil velocity is reduced in T1 transitions in *Src42A^i^* embryos (Fig. 5D). From these experiments, we conclude that Src42A is involved in generating or maintaining tension in intercalating cells at T1-transitions.

**Figure 5:**
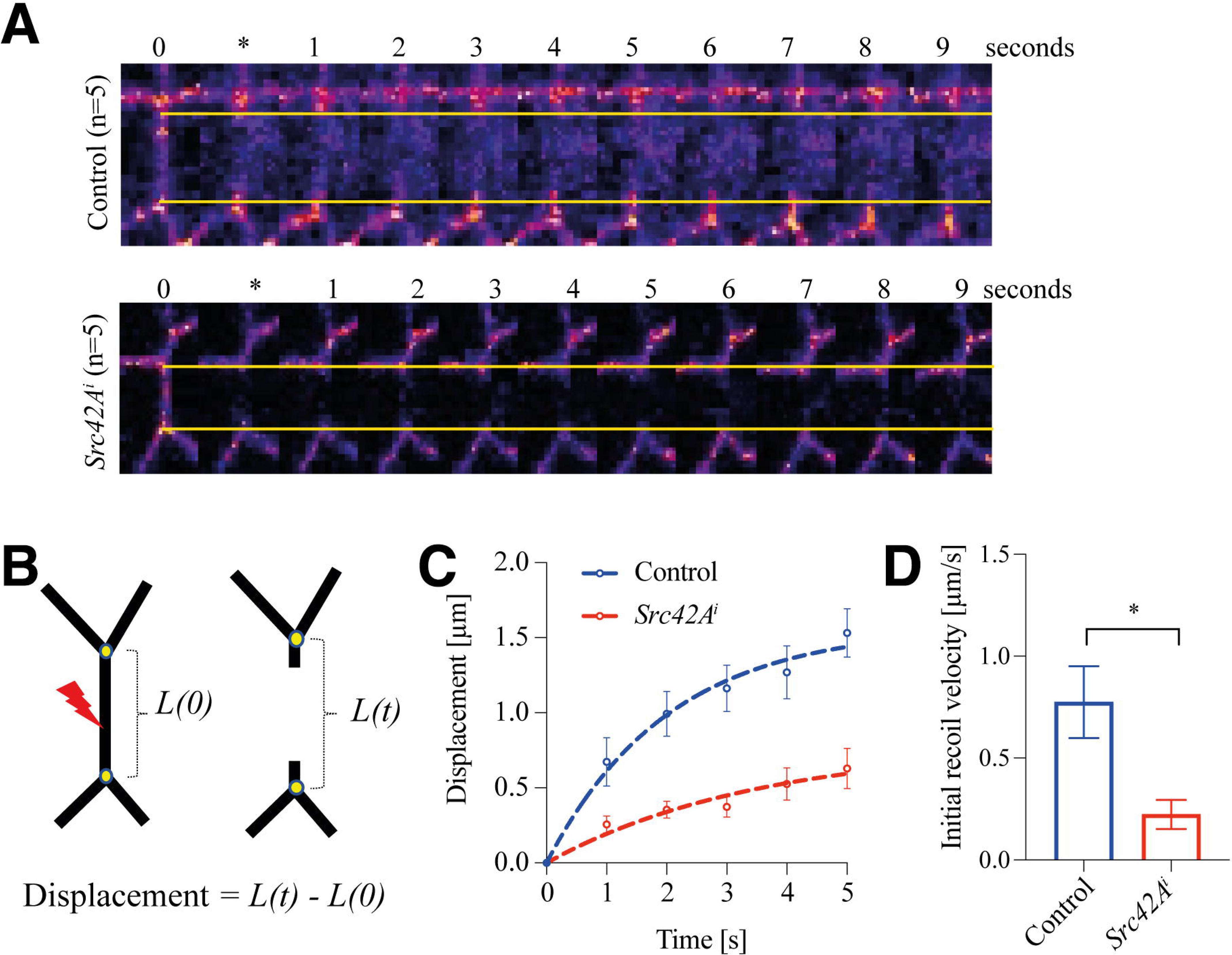
Src42A alters tension at the AP junction border. **(A)** Laser ablation experiment conducted in control and *Src42A^i^* embryos, length between the tAJs is marked in yellow line, movies were recorded for every one second after ablation of a AP junction border (time scale in seconds at the top of images; * time point at which laser cut was performed; n = embryos examined). **(B)** Schematic view of laser ablation experiment and displacement calculation. Yellow dots represent tAJs, *L(0)* is the initial length between vertices, *L(t)* is the length between vertices over time. **(C)** tAJs displacements were plotted for control and *Src42A^i^*, displacement values are marked in small circles with respective error values, non-linear regression is marked in dotted lines. **(D)** Initial recoil velocity was calculated from the slope of non-linear regression curve showing significant difference between control and *Src42A^i^*. Unpaired t-test shows p value of 0.0165 (less than 0.05).

### Src42A acts together with Abl in controlling E-cadherin and β-catenin levels at the AP cell interfaces

In *Drosophila*, Src42A genetically interacts with components of the E-cadherin/β-catenin complex and Src42A can be detected in a ternary complex with E-cadherin and β-catenin (Takahashi et al. 2005). Furthermore the E-cadherin-catenin complex, which is essential for cell-cell adhesion, is regulated by phosphorylation of Y654 of β-catenin by mammalian c-src (Röper et al. 2018). β-catenin is preferentially localized at the DV cell border during planar polarization of germband cells and the phosphorylation of β-catenin Y667 by Abelson kinase (Abl) is vital to achieve this polarization (Tamada et al. 2012). It was also shown in the *Drosophila* wing epithelium that Src42A can potentially activate Abl kinase (Singh et al. 2010). These data raise the hypothesis that Src42A affects the E-cadherin/ß-catenin complex indirectly via Abl. In *Src42Ai* embryos, there was no significant change in protein levels at the DV cell border, in contrast the AP borders show enhanced levels of β-catenin and E-cadherin (Fig. 6A,B,C). These data indicate that the planar polarity of the E-cadherin-catenin complex is compromised in *Src42A^i^* embryos. To decide whether Src42A acts in the same or in distinct pathways with respect to Abl, we conducted a double knockdown experiment targeting both kinases. Germband elongation was examined by time-lapse video microscopy (Fig. 7A). The cumulative displacement values were plotted over time for control and *Src42A^i^*+*Abl^i^* and the speed of elongation was calculated from the displacement data for the double knockdown in comparison with *Src42A^i^* (Fig. 7B,C). We find that in fast phase *Src42A^i^* and *Src42A^i^+Abl^i^* delay germband extension to a similar degree. However, during the slow phase of extension *Src42A^i^+Abl^i^* showed much stronger delay compared to *Src42A^i^* alone (Fig. 7D,E). Furthermore, the speed was calculated from the displacement data for the double knockdown in comparison with *Src42A^i^* (Fig. 7C). While there was no difference in the fast phase, in slow phase the speed was more strongly reduced in *Src42Ai*+*Abl^i^* compared to *Src42A^i^* alone (Fig. 7D,E). Therefore, *Src42A* and *abl* may function in parallel pathways during the slow phase of germband extension.

**Figure 6:**
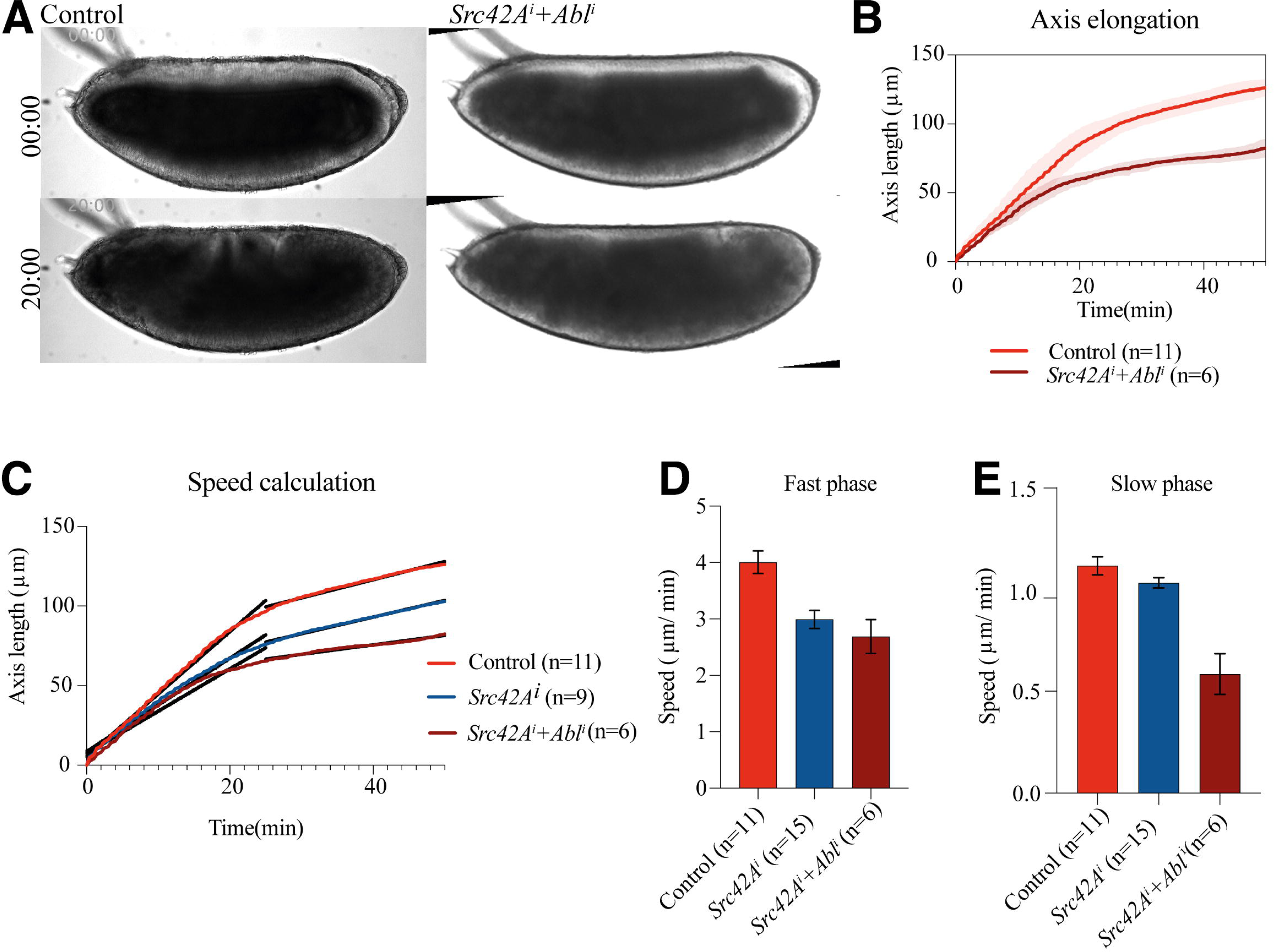
Planar polarized distribution of β-catenin and E-cadherin. **(A)** Control and *Src42A^i^* fixed embryos showing the apical area of germband cells stained for β-catenin (β-Cat), E-cadherin (ECad) and Src42A (Src42A and His-GFP was imaged on same channel). Scale bar represents 10µm. **(B,C)** Quantification of β-Cat and Ecad at the DV border and AP border of bAJs. Immunostaining intensities were measured in at least 5 bAJs for each embryo (indicated as black dots in the graphs), n value denotes the number of embryos analysed. Nested t-test were performed on all data p values for β-Cat at DV border and AP border are 0.4236 and 0.0037 respectively, p values for ECad at DV border and AP border are 0.9700 and 0.0499 respectively.

**Figure 7:**
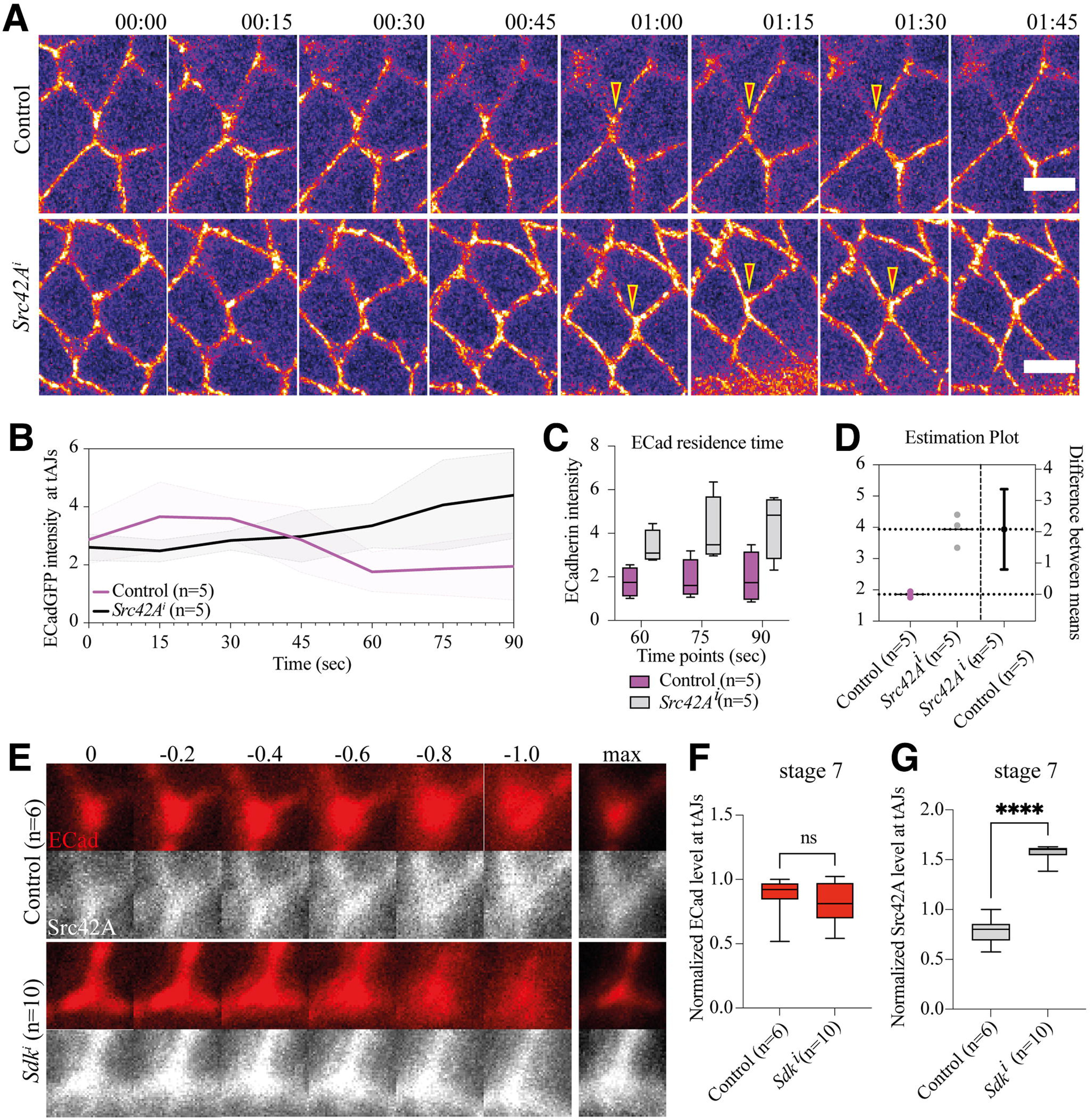
Src42A and Abelson double knockdown enhances germband extension delay. **(A)** Bright field images of control and *Src42A^i^+Abl^i^* shows delay in germ-band extension. Time series is mentioned on the left side of the images in mm:ss format. **(B)** Cumulative displacement of germband cells is plotted over time. **(C,D**,**E)** Slope was calculated for slow and fast phase of germ-band extension in control, *Src42A^i^* and *Src42A^i^+Abl^i^* (slope values demonstrate the speed differences between the respective genotypes).

### Src42A controls E-cadherin turnover at tAJs in T1 transition cells

We found that Src42A showed a distinct localization at the tAJs (Fig. 1B). The tAJs is one of the tension hotspots during germband elongation, and Sidekick (Sdk) protein is a core component of this subcellular domain (Uechi and Kuranaga 2019; Finegan et al. 2019; Salomon et al. 2017). Sdk also regulates E-cadherin endocytosis during cell intercalation when new junction formation occurs at the horizontal interface (Letizia et al. 2019). Despite the requirement of Sdk, the mechanisms controlling E-cadherin dynamics at the tAJs is not well understood. Therefore, we recorded E-cadherin-GFP localization in the background of *Src42A^i^* during T1 transition. The localization of E-cadherin-GFP exhibits an accumulation and dispersion cycle during T1 transition at tAJs (Fig. 8A) (Vanderleest et al. 2018). In *Src42A^i^* embryos E-cadherin levels were continuously increasing at the tAJs suggesting that the turnover of E-cadherin is affected during T1 transition. From the vertex intensity ratio calculation, E-cadherin levels are not fluctuating in embryos deficient for Src42A, the estimation plot showing the significant difference among the mean E-cadherin values for control and *Src42A^i^* embryos (Fig. 8B,C,D). Further, stage 7 *Sdk^i^* embryos were subjected to immunostaining experiments to check E-cadherin and Src42A localization (Fig. 8E), and protein levels at the tAJs were measured from maximum intensity projection images. The result show that there were no change in E-cadherin level, however Src42A level at the tAJs were increased (Fig. 8F,G). We conclude that E-cadherin turnover is affected at tAJs when Src42A was knocked-down.

**Figure 8:**
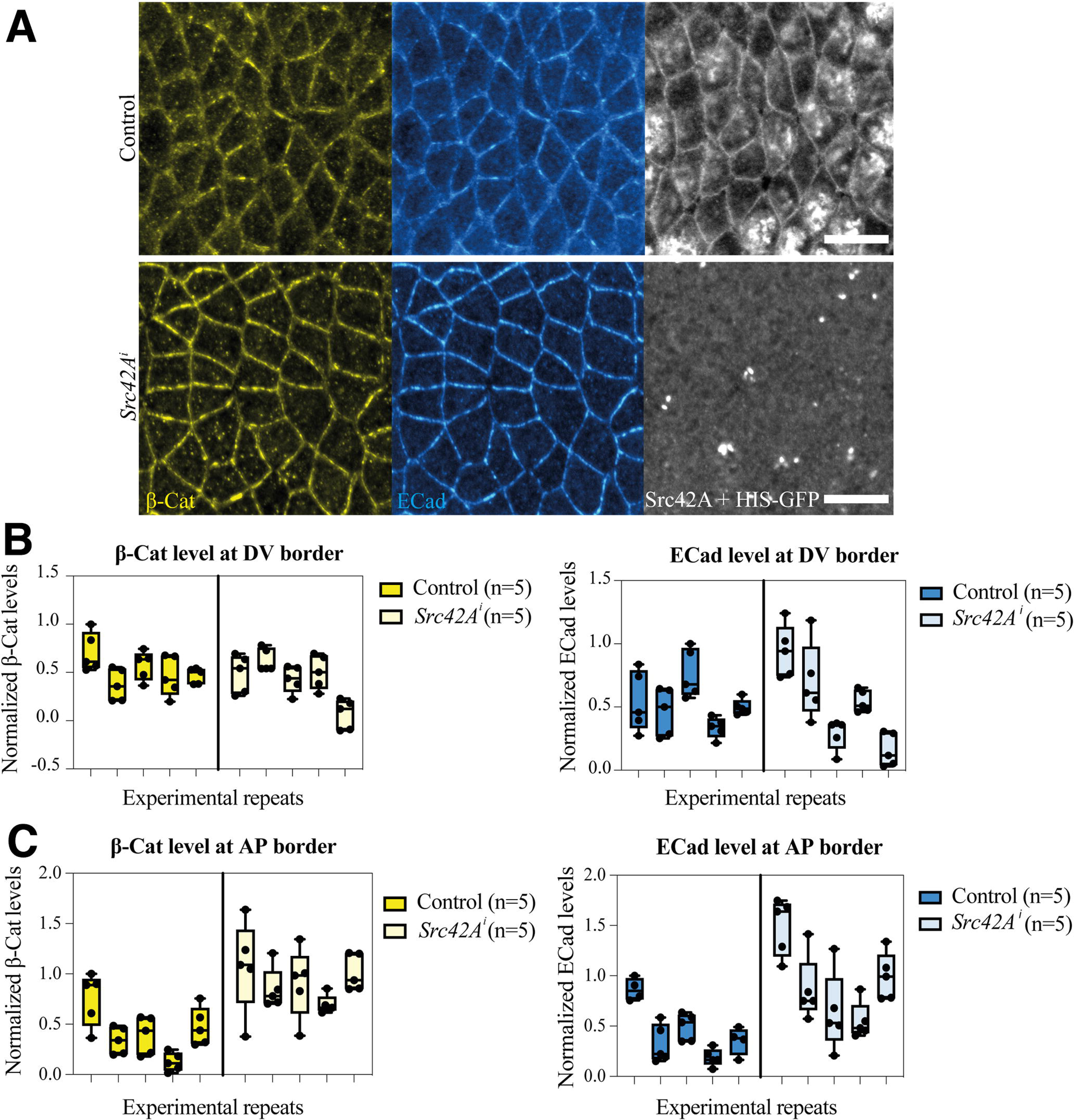
Src42A controls E-cadherin intensity at tAJs during T1-transition. **(A)** Maximum intensity projection of 5 individual images taken as Z-stacks with an interval of 0.3µm in between each other. Control and *Src42A^i^* show E-cadherin levels during T1 transition were analysed using maximum intensity projection images (Time indicated at the top of images in mm:ss format; arrowheads indicates tAJs and scale bars corresponds to 5µm). **(B)** E-cadherin (Ecad) intensity at tAJs were normalized to bAJs intensity and background and plotted over time, control (magenta) show fluctuation whereas *Src42A^i^* show increasing levels of ECad. **(C)** The intensity at the last three timepoints plotted between control and *Src42A^i^* and show a significant difference in the mean values in the estimation plot **(D)**. Unpaired t-test was performed with Welsch’s correction (p value = 0.0190) indicating significant difference between control and *Src42A^i^*. **(E)** Control and *Sdk^i^* embryos imaged at stage 7, the vertex intensity of Ecad and Src42A imaged in different Z-stack level with 0.2µm interval from the surface (shown at the top of the images), max denotes the maximum intensity projection obtained from consecutive Z-stacks and n value represents the number of embryos. **(F,G)** E-cadherin level show no significant (p = 0.1746) difference between the control and *Sdk^i^* whereas Src42A level at the tAJs is increased significantly (p = 0.0001).

### Src42A germline clone analysis confirms the *Src42A^i^* phenotypes

The use of transgenic RNAi to knockdown maternal Src42A has caveats including off-target effects and incomplete depletion due to perdurance of maternal Src42A gene product. Attempts to confirm the specificity of the RNAi knock-down by rescuing *Src42A^i^* embryos using Src42A transgenes were inconclusive, because Src42A overexpression, in particular of tagged versions, can induce lethality (Figure S3) (Irby and Yeatman 2000). Therefore, to confirm the specificity of the effects caused by RNAi knockdown experiments, we generated germline clones by inducing double strand breaks at a chromosomal site (*stlk*; 41A3) proximal to the *Src42A* (42A6-7) locus using CRISPR-Cas9 (Allen et al. 2021). In *Src42A^26-1^* germline clones (named *Src42A^GLC^* in this study), the cytoplasmic clearing is consistently defective during cell cycle 13 to 14 transition; interestingly the phenotype appears much stronger as the furrows looks uneven compared to *Src42A^i^* pointing to a yet unknown function of Src42A in cellularization (Fig. 9A & Fig. 2B). The ovaries from *Src42A^GLC^* were dissected to check the level of Src42A in germline cells and found that Src42A is completely absent in nurse cells. The absence of Src42A in the germline also affected the localization of E-cadherin at the nurse cell boundaries (Fig. 9B). We further analyzed the speed of germband elongation (Fig. 9 C,D) and the planar polarized distribution of β-catenin and E-cadherin (Fig. 9 E-I). The phenotype of *Src42A^GLC^* was stronger compared to *Src42A^i^*, suggesting that some Src42A maternal gene products escape the RNAi knockdown. This experiment provides evidence that the knockdown effects in *Src42A^i^* is specific to reduction of Src42A and indicates that Src42A is also involved in normal germline development and blastoderm development.

**Figure 9:**
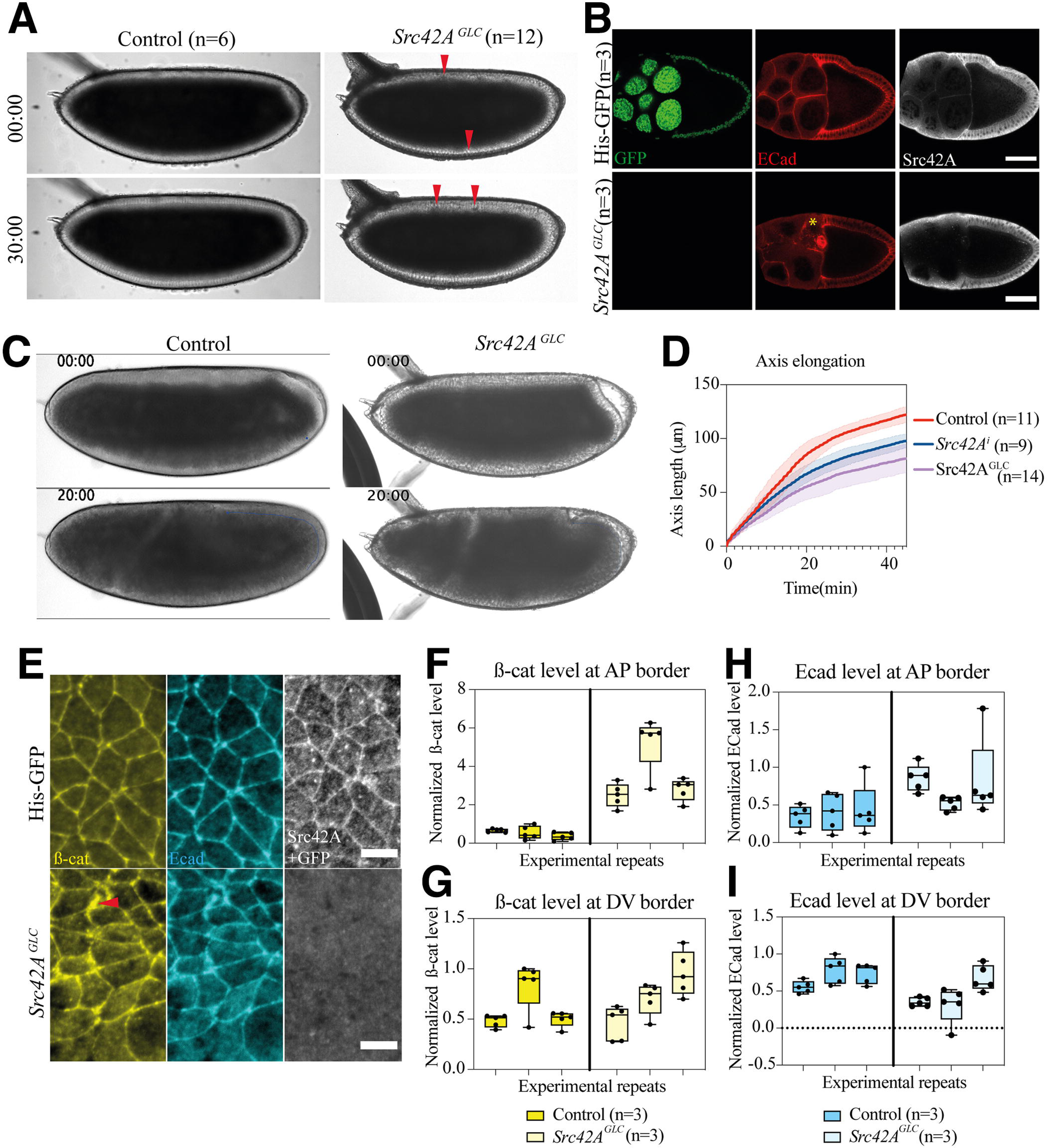
Phenotypic analyses of Src42A germline clones. **(A)** Bright field images of control and *Src42A^GLC^* embryos showing defective yolk cytoplasm (marker in red arrow heads). **(B)** Stage 9 control and *Src42A^GLC^* oocytes stained for Ecad and Src42A, asterisk marking the Ecad distribution difference (scale bar denotes 50 μm). **(C)** Images taken from bright field movies of control and *Src42A^GLC^* embryos showing a delay in germband extension. **(D)** Graph showing defect in germ-band axis elongation in *Src42A^GLC^* in comparison to control and *Src42A^i^* conditions. **(E)** Control and *Src42A^GLC^* fixed embryos showing β-cat and E-cadherin distribution in stage 7 embryos (red arrow heads mark the defects in β-cat distribution, scale bar denotes 5 μm). **(F,G,H,I)** Normalized at and Ecad level at the AP and DV border of the cells were measured and plotted between control and *Src42A^GLC^* embryos. Nested t-test were performed on all data p values for β-Cat at DV border and AP border are 0.6183 and 0.0268 respectively, p values for ECad at DV border and AP border are 0.1380 and 0.0370 respectively.

## Discussion

In this study we aimed to understand the function of Src42A in regulating the dynamics of E-cadherin at the bAJs and tAJs during germband extension. Depletion of maternal and zygotic Src42A by RNAi knock-down resulted in defects in early embryogenesis, most notably an impaired planar polarity of E-cadherin and β-catenin. In particular the E-cadherin intensity at AP cell border was abnormally high, suggesting that E-cadherin turnover at the AP border is affected. During this stage, E-cadherin polarity was shown to control the flow of actomyosin to the AP cell border (Levayer and Lecuit 2013). In the absence of E-cadherin endocytosis, the MyoII flow required for vertical junction constriction may be compromised. This conclusion is consistent with the reduced Myo II polarity (Tamada et al., 2021) and the reduced recoil velocity that we observed in *Src42A^i^* embryos when the AP border is photo ablated. Less tension at the AP border means that the constriction rate of vertical AJs at the T1-transition is compromised. Our findings support the evidence that Src kinases (both Src42A and Src64B) act downstream of Toll-2 and Toll-6 receptors and thereby alter formation of Myo II cables at the AP cell border and Bazooka localization at the DV cell border (Tamada et al. 2021). Therefore, knocking down Src42A alters the dynamics of molecular components in T1 transition cells and thereby compromises the speed of germband extension. We also generated for the first time germline clones for a mutant Src42A allele using a CRISPR/Cas9 approach that can be used to generate germline or somatic mosaic clones for genes which are present proximal to the FRT site in the centromeric region. The embryos derived from Src42A germline clones recapitulates the phenotypes of Src42A knockdown experiments and extends the spectrum of requirements of maternal Src42A for oogenesis and blastoderm formation.

Previous studies on Abl kinase show that it can alter the planar polarity of β-catenin during rosette formation (Tamada et al. 2012). During cell invasion, Src can act upstream of Abl and there is a potential feedback from Abl which has a positive impact on Src42A (Singh et al. 2010). Despite the interplay between Src and Abl kinases, the relationship between these two kinases was not studied during germband elongation. Here we show that RNAi knockdown of both, Src42A and Abl kinases, resulted in an enhanced phenotype during the slow phase of germband extension. While the molecular details of this genetic interaction remain to be addressed, we conclude that Src42A and Abl act in parallel genetic pathways during the slow phase where the rosettes resolve by extending new horizontal junctions.

Abelson kinase (Abl) is required for the mobility of Canoe (*Drosophila* homolog of Afadin) from tAJs to bAJs and this mobility depends on tyrosine phosphorylation of Canoe by Abl as a Canoe phosphomutant displayed delayed T1 transition (Yu and Zallen 2020). Canoe maternal mutants show similar defects like *Src42A^i^* embryos, both display less phosphotyrosine signal at the tAJs. Src42A can also phosphorylate Smallish which required for the planar polarity of Baz and Canoe at the bAJs and localization of Smallish is enriched at tAJs during germband extension stages (Beati et al. 2018). It is known that Canoe acts as linker between the junctional cadherin-catenin complex and the actin cytoskeleton. We hypothesized that Src42A can activate Abl in mobilizing Canoe therefore, we generated embryos with double knockdown for Src42A and Abl where the slow phase of germ-band extension is delayed compared to single Src42A knockdown. When Abl alone was knocked down Canoe showed impaired mobility at the AJs (Yu and Zallen 2020). If Src42A acts directly on Canoe or indirectly through Abl in mobilizing Canoe between bAJs and tAJs, this would result in highly disproportionate Canoe localization at both tAJs and bAJs in double knockdown. In mammals, Afadin regulates E-cadherin turnover at cell junctions by trans-interacting with Nectin, if there is no trans interaction, E-cadherin is internalized by Clathrin-mediated endocytosis (de Beco et al. 2009; Takeichi 2014; Hoshino et al. 2005). In our analysis we find that E-cadherin intensity at tAJs remains high in *Src42A^i^* embryos compared to wild-type embryos. This implicates that more E-cadherin is accumulating at the tAJs, which could possibly occur due to reduced levels of Canoe causing more E-cadherin clusters at the tAJs during T1-transition.

At mid stage of germband extension, E-cadherin protein level at the tAJs exhibits an accumulation and dispersion cycle that correlates with cell intercalation (Vanderleest et al. 2018). The resident tricellular junction protein Sdk regulates E-cadherin endocytosis during genitalia rotation, where cell intercalation occurs continuously (Uechi and Kuranaga 2019). Sdk connects to the actin cytoskeleton through Canoe and Polychaetoid which transmits tension at the tAJs (Letizia et al. 2019). In *Src42A^i^* embryos, E-cadherin intensity is continuously increased at tAJs during cell intercalation which suggest a specific role for Src42A in maintaining E-cadherin turnover in tAJs. What acts upstream of Src42A at tAJs in mediating E-cadherin turnover remains elusive. We aimed to understand the relationship between Sdk and Src42A in context to E-cadherin turnover. In *Sdk^i^* conditions, we were not able to observe any difference in E-cadherin intensity in fixed embryos but the level of Src42A is increased at tAJs. These data suggest that Sdk directly or indirectly controls the recruitment of Src42A to the tAJs, but the exact relationship between Sdk, Src42A and E-cadherin turnover remains to be addressed.

In this study, we provide evidence for dynamic function of Src42A in detail on how it is regulating molecular components at the bAJs and tAJs. One of the interesting aspects of this study is during T1 transition Src42A regulates the residence time of E-cadherin which points out that Src42A can have potential function at the tAJs which has not been reported so far. The mechanical tension at the tAJs can potentially transduced into the cell via Src42A. In addition to the involvement in planar cell polarity Src42A may have additional function at the tAJs in regulating E-cadherin turnover. In this context it would be interesting to identify tAJs resident proteins which act upstream and downstream of Src42A.

## Materials and methods

### Molecular biology and antibody generation

Full length *Src42A* cDNA was cloned into *pGGWA* vector using the Gateway cloning method (Katzen 2007) for antibody generation. The final *pGGWA+Src42A* vector contains a GST (Glutathione S-transferase) tag at the N-terminal end of *Src42A*, further the recombinant protein *GST-Src42A* was expressed in *BL21*(DE3) *E*. *coli* cells at 18°C for overnight and purified using affinity chromatography (using Glutathione Sepharose beads). Guinea pigs were used for antibody generation (Eurogentec, Belgium). After immunization, the serum from the final bleed was directly used as Src42A primary antibody with a dilution of 1:500 for all experiments.

### Fly genetics

All fly stocks were raised at 25°C. To analyse the dynamic localization of Src42A during cellularization and germband extension, embryos from *Sqh::KI-GFP* flies were fixed and stained. Additionally, *Src42A^26-1^* flies (zygotic mutant) (Takahashi et al. 2005) were used to study the specificity of the generated antibody. To check the cross-reactivity of Src42A antibody with Src64B, *UASp*>*Src42A*-*HA* and *UASp>Src64B-HA* female flies (gift from Andreas Wodarz, University of Cologne, Germany) were crossed with *engrailed*>*Gal4* male flies.

Maternal and zygotic knock-down of Src42A was performed using the *UAS*/*GAL4* system (Staller et al. 2013; Brand and Perrimon 1993). Female flies carrying *shRNA* for *Src42A* under the control of *UAS* promoter (y^1^ sc* v^1^ sev^21^; P{TRiP.HMC04138}attP2/TM3, Sb^1^) were crossed to male driver flies (y^1^ w*; P{mat 4-GAL4-VP16}67; P{mat 4-GAL4-VP16}15). In the next generation P{mat 4-GAL4-VP16}67/ +; P{mat 4-GAL4-VP16}15/ P{TRiP.HMC04138}attP2 flies were collected and both males and females from these genotypes were crossed to obtain embryos in which maternal and zygotic Src42A (named *Src42A^i^* in this work) was knocked down in the F2 generation. As controls in germ-band elongation assays, male driver lines (y^1^ w*; P{mat 4-GAL4-VP16}67; P{mat 4-GAL4-VP16}15) were crossed to w^1118^ females, the F1 generation was backcrossed and the F2 embryos were analysed. For all immunostaining quantification experiments *His-GFP* embryos were used as a control and stained together in a same tube along with *Src42A^i^* embryos. For dynamic analysis of T1 transitions, y^1^ w*; P{mat 4-GAL4-VP16}67 sqh::Utrophin-GFP; P{mat 4-GAL4-VP16}15/TM6 flies were used as a driver to obtain *Src42A^i^* embryos; *Sqh::Utrophin GFP* flies were used as control.

For E-cadherin vertex intensity analysis and laser ablation experiments y^1^ w*; P{mat 4-GAL4-VP16}67 Shg::DECadGFP/ Shg::DECadGFP; P{mat 4-GAL4-VP16}15/ P{TRiP.HMC04138}attP2 flies were crossed to obtain *Src42^i^* embryos along with E-cadherin marker; Shg::DECadGFP flies were used as control. For Src42A and Abelson double knockdown (*Src42A^i^*_*Abl^i^*) P{TRiP.HMC05140}attP40; P{TRiP.HMC04138}attP2/TM3, Sb^1^ females were used to cross with same maternal driver line as mentioned above. RNAi experiments were also performed for *sidekick* (*sdk*) using y^1^ sc* v^1^ sev^21^; P{TRiP.HMS00292}attP2 fly line (mentioned as *Sdk^i^*). All the TRiP fly lines were generated at Harvard Medical School for the Transgenic RNAi Project (Ni et al. 2011).

Germ-line clones lacking Src42A were generated using a CRISPR-Cas9 approach based on gRNA-induced double-strand (ds) breaks at a site (*stlk* locus; 41A3) proximal to the *Src42A* (42A6-7) locus. A transgene constitutively expressing *stlk* gRNA *(P{TKO.GS00956}attP40(y^+^)*; Bloomington #76505) was recombined with *Src42A^26-1^*. To induce ds breaks in the non-essential *stlk* gene, *y w; stlk-gRNA-attP40(y^+^) Src42A^26-1^* females were crossed with *y w act5c-Cas9 lig4/Y; FRT42D ovoD(w^+^)/+* males. All eggs produced by F1 females of the genotype *y w act5c-Cas9 lig4 / y w; stlk-gRNA-attP40(y^+^) Src42A^26-1^* / *FRT42D ovoD(w^+^)* are derived from germline cells lacking the dominant female-sterile *ovoD* transgene (Chou and Perrimon 1996) and are therefore homozygous for *Src42A^26-1^*. Control females carrying the *FRT42D ovoD(w^+^)* chromosome in the absence of the *act5c-Cas9* source or of the *stlk-gRNA* source did not lay any eggs. Females carrying *Src42A^26-1^* germline clones were crossed with *w*/Y; Src42A^26-1^/ CyO[twi::GFP]* males to obtain embryos lacking maternal and zygotic *Src42A*. The genotypes of the germline clone embryos were identified by no *twi::GFP* expression.

### Immunostainings and Immunoblotting

Embryos were collected from 0 to 7 hours and fixed using 4% formaldehyde made in phosphate buffer saline as described (Müller 2008). After fixation the embryos were blocked in 5% BSA and stained with the following primary antibodies guineapig anti-Src42A (1:500), rabbit anti-Bazooka (1:2000) (Wodarz et al. 1999), mouse anti-phosphotyrosine20 (1:1000) (purchased from BD Biosciences), rabbit anti-beta galactosidase (1:1000) (purchased from Cappel Research Reagents), rat anti-E-cadherin (1:20) (Developmental Studies Hybridoma Bank), mouse anti-Arm^N2^ (1:250) (Developmental Studies Hybridoma Bank) and rat anti-haemagglutinin (1:1000) (Roche). The following secondary antibodies were used, goat anti-guineapig Alexa 647 (Invitrogen), donkey anti-rabbit Cy3 (Stratech), donkey anti-mouse Cy3 (Jackson Immuno Research labs), donkey anti-rat Cy2 and donkey anti-rat Cy3 (Stratech). All secondary antibodies were used at 1:250 dilution.

For immunoblotting experiment, staged embryos were collected, and lysates were prepared under denaturing conditions using 2x SDS sample buffer (0.125M Tris with pH 6.8, 20% Glycerin, 4% SDS, 0.004% bromophenolblue and 10% beta-mercaptoethanol). To check the knockdown efficiency, embryos were collected from 0 to 7 hours, protein lysates were prepared using RIPA buffer (50mM Tris, 150mM NaCl, 1% NP40, 0.1% SDS, 0.5% Na-Deoxycholate and 1%Triton X-100) under non-denaturing conditions and separated on a 15% SDS-PAGE gel (Wodarz 2008), transferred onto nitrocellulose membrane (Amersham™ Protron™ 0.2µm NC) and the membrane is blocked and stained with primary antibodies Guineapig anti-Src42A (1:500), rabbit anti-haemagglutinin (1:1000) (Sigma-Aldrich) and mouse anti-alpha tubulin (1:2000) (Developmental Studies Hybridoma Bank) and the following secondary antibodies donkey anti-guinea pig IR dye 800 (1:10000) (LI-COR Biosciences), donkey anti-mouse IR dye 680 (1:10000) (LI-COR Biosciences) and donkey anti-mouse HRP (1:2000) (Roche) were used to detect the protein levels in LI-COR Odyssey^®^ Fc imaging system.

### Image acquisition and quantification

#### Protein levels at bicellular and tricellular AJs

Images were acquired using confocal laser scanning microscope (Zeiss LSM880) for 1.2 µm depth from the surface in Z-plane with 0.2µm interval. All the images were processed and analysed using Fiji. In total, 6 images were acquired, and maximum intensity projection (MIP) images were generated from these images. Using the MIP images quantifications were performed. Average pixel intensity at the two and three cell contacts were measured and subtracted with the background value. The pixel intensities were normalized to the maximum value of control image. Both control and *Src42Ai* embryos were stained in the same tube for quantification purpose.

#### Germ-band elongation assay

Bright field images were taken from Zeiss Axiophot and Olympus BX61 microscope and processed using Fiji. The length of germ-band elongation was tracked using ‘manual tracking’ plugin, and the cumulative displacement of germ-band were plotted over time. To access the speed of germ-band elongation a linear curve was drawn for slow and fast phase of germ-band elongation, using linear curve slope was calculated for respective phases.

#### T1 transition analysis

Live imaging experiments were performed using sqh::Utrophin-GFP as a marker for cell membrane. Imaging was performed in confocal laser scanning microscope (LSM880) in airy scan mode. The transition between stage 6 and 7 were recorded with 15 secs time interval. All the acquired images were analysed using ‘Tissue Analyzer’ plugin in Fiji. Images from each timepoints were segmented and the length of AP axis and DV axis was measured from segmented images. Initial AP axis length (L0) is normalized to the length over time (Lt) and the normalized value was plotted over time.

#### E-cadherin vertex intensity ratio

Images were taken in confocal laser scanning microscope in airy scan mode. Germ-band cells undergoing T1 transitions were imaged for a depth of 1µm in Z-plane with 0.2µm interval. MIP images were generated from the acquired images for all time points. Cells undergoing T1 transition containing 2.5µm AP axis length were selected and used for analysis. From the final images E-cadherin vertex intensity ratio is calculated using the following formula (ItAJs-IB)/(IbAJs-IB) (Vanderleest et al. 2018). The vertex intensity ratio was then plotted over time.

### Laser ablation

Stage 7 embryos expressing ECadGFP were prepared for live imaging and recorded in the GFP channel with 1sec time interval on a confocal laser scanning microscopy (Zeiss, LSM980 with 100x magnification using oil immersion, 1.4 NA). A 355 nm pulsed laser (DPSL355/14, 355 nm, 70 µJ/pulse, Rapp OptoElectronic) was employed for ablation and manipulated on the ‘REO-SysCon-Zen’ platform (Rapp OptoElectronic). The 355 nm pulsed laser was mounted on epiport of the confocal laser scanning microscopy. Laser ablation was performed with 5% of laser power, with 200 ms (around 40 pules) exposure time at the AP axis of the cells undergoing T1 transition. For analysis, the displacement length of ablated tAJs (L(t)) were measured manually in Fiji. The displacement value were normalized to the initial length (L(0)) between tAJs and plotted over time. The initial recoil velocity were calculated using Kelvin-Voigt fibber model to the following equations on Prism8 (Liang et al. 2016; Fernandez-Gonzalez et al. 2009).

Extraction of initial recoil velocity value:

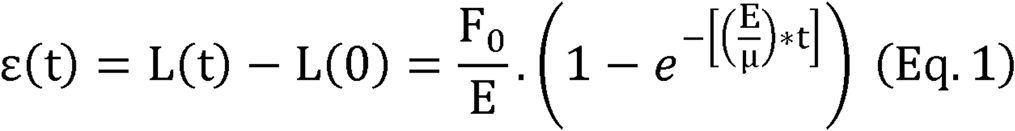

were,

F_0_ is the tensile force present at the junction before ablation,

E is the elasticity of the junction,

μ is the viscosity coefficient related to the viscous drag of the cell cytoplasm.

As fitting parameters for the above equation, we introduced

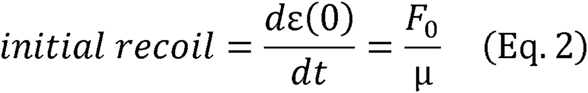

## Acknowledgements

We thank Andreas Wodarz for providing fly stocks and antibodies. We thank Tanja Wilhelm, Christine Otto, Nicole Schleinschok for fly maintenance and expert technical assistance, and Monika Winneknecht and Birgit Simon for preparing fly media. We thank the Bloomington Drosophila stock centre (Bloomington, USA) for Drosophila stocks, the Developmental Studies Hybridoma Bank (Iowa, USA) for antibodies and the Transgenic RNAi Project (Harvard University, USA) for providing RNAi lines. Work in SL’s laboratory was supported by the Deutsche Forschungsgemeinschaft (SFB 1348 “Dynamic Cellular Interfaces”; SFB 1009 “Breaking Barriers”), the “Cells-in-Motion” Cluster of Excellence (EXC 1003-CiM) at the University of Münster, and the University of Münster. Work in HAM’s laboratory was supported by a new investigator award to HB and core funding from the University of Kassel and the PhosMOrg Consortium (Univ. Kassel).

## Notes

### Competing Interest Statement

The authors have declared no competing interest.

## References

Allen, S.E., Koreman, G.T., Sarkar, A., Wang, B., Wolfner, M.F. and Han, C. 2021. Versatile CRISPR/Cas9-mediated mosaic analysis by gRNA-induced crossing-over for unmodified genomes. PLoS Biology 19(1), p. e3001061.

Ambrosini, A., Rayer, M., Monier, B. and Suzanne, M. 2019. Mechanical Function of the Nucleus in Force Generation during Epithelial Morphogenesis. Developmental Cell 50(2), p. 197–211.e5.

Beati, H., Peek, I., Hordowska, P., et al. 2018. The adherens junction-associated LIM domain protein Smallish regulates epithelial morphogenesis. The Journal of Cell Biology 217(3), pp. 1079–1095.

de Beco, S., Gueudry, C., Amblard, F. and Coscoy, S. 2009. Endocytosis is required for E-cadherin redistribution at mature adherens junctions. Proceedings of the National Academy of Sciences of the United States of America 106(17), pp. 7010–7015.

Bertet, C., Sulak, L. and Lecuit, T. 2004. Myosin-dependent junction remodelling controls planar cell intercalation and axis elongation. Nature 429(6992), pp. 667– 671.

Blankenship, J.T., Backovic, S.T., Sanny, J.S.P., Weitz, O. and Zallen, J.A. 2006. Multicellular rosette formation links planar cell polarity to tissue morphogenesis. Developmental Cell 11(4), pp. 459–470.

Brand, A.H. and Perrimon, N. 1993. Targeted gene expression as a means of altering cell fates and generating dominant phenotypes. Development 118(2), pp. 401–415.

Brunet, T., Bouclet, A., Ahmadi, P., et al. 2013. Evolutionary conservation of early mesoderm specification by mechanotransduction in Bilateria. Nature Communications 4, p. 2821.

Chou, T.B. and Perrimon, N. 1996. The autosomal FLP-DFS technique for generating germline mosaics in Drosophila melanogaster. Genetics 144(4), pp. 1673–1679.

Fernandez-Gonzalez, R., Simoes, S. de M., Röper, J.-C., Eaton, S. and Zallen, J.A. 2009. Myosin II dynamics are regulated by tension in intercalating cells. Developmental Cell 17(5), pp. 736–743.

Finegan, T.M., Hervieux, N., Nestor-Bergmann, A., Fletcher, A.G., Blanchard, G.B. and Sanson, B. 2019. The tricellular vertex-specific adhesion molecule Sidekick facilitates polarised cell intercalation during Drosophila axis extension. PLoS Biology 17(12), p. e3000522.

Gheisari, E., Aakhte, M. and Müller, H.-A.J. 2020. Gastrulation in Drosophila melanogaster: Genetic control, cellular basis and biomechanics. Mechanisms of Development 163, p. 103629.

Hoshino, T., Sakisaka, T., Baba, T., Yamada, T., Kimura, T. and Takai, Y. 2005. Regulation of E-cadherin endocytosis by nectin through afadin, Rap1, and p120ctn. The Journal of Biological Chemistry 280(25), pp. 24095–24103.

Irby, R.B. and Yeatman, T.J. 2000. Role of Src expression and activation in human cancer. Oncogene 19(49), pp. 5636–5642.

Katzen, F. 2007. Gateway(®) recombinational cloning: a biological operating system. Expert opinion on drug discovery 2(4), pp. 571–589.

Kong, D., Wolf, F. and Großhans, J. 2017. Forces directing germ-band extension in Drosophila embryos. Mechanisms of Development 144(Pt A), pp. 11–22.

Letizia, A., He, D., Astigarraga, S., et al. 2019. Sidekick Is a Key Component of Tricellular Adherens Junctions that Acts to Resolve Cell Rearrangements. Developmental Cell 50(3), p. 313–326.e5.

Levayer, R. and Lecuit, T. 2013. Oscillation and polarity of E-cadherin asymmetries control actomyosin flow patterns during morphogenesis. Developmental Cell 26(2), pp. 162–175.

Levayer, R., Pelissier-Monier, A. and Lecuit, T. 2011. Spatial regulation of Dia and Myosin-II by RhoGEF2 controls initiation of E-cadherin endocytosis during epithelial morphogenesis. Nature Cell Biology 13(5), pp. 529–540.

Liang, X., Michael, M. and Gomez, G.A. 2016. Measurement of Mechanical Tension at Cell-cell Junctions Using Two-photon Laser Ablation. Bio-protocol 6(24).

Müller, H.-A.J. 2008. Immunolabeling of embryos. Methods in Molecular Biology 420, pp. 207–218.

Ni, J.-Q., Zhou, R., Czech, B., et al. 2011. A genome-scale shRNA resource for transgenic RNAi in Drosophila. Nature Methods 8(5), pp. 405–407.

Paré, A.C., Vichas, A., Fincher, C.T., et al. 2014. A positional Toll receptor code directs convergent extension in Drosophila. Nature 515(7528), pp. 523–527.

Paré, A.C. and Zallen, J.A. 2020. Cellular, molecular, and biophysical control of epithelial cell intercalation. Current Topics in Developmental Biology 136, pp. 167– 193.

Rauzi, M., Lenne, P.-F. and Lecuit, T. 2010. Planar polarized actomyosin contractile flows control epithelial junction remodelling. Nature 468(7327), pp. 1110–1114.

Röper, J.-C., Mitrossilis, D., Stirnemann, G., et al. 2018. The major β-catenin/E-cadherin junctional binding site is a primary molecular mechano-transductor of differentiation in vivo. eLife 7.

Salomon, J., Gaston, C., Magescas, J., et al. 2017. Contractile forces at tricellular contacts modulate epithelial organization and monolayer integrity. Nature Communications 8, p. 13998.

Singh, J., Aaronson, S.A. and Mlodzik, M. 2010. Drosophila Abelson kinase mediates cell invasion and proliferation through two distinct MAPK pathways. Oncogene 29(28), pp. 4033–4045.

Staller, M.V., Yan, D., Randklev, S., et al. 2013. Depleting gene activities in early Drosophila embryos with the “maternal-Gal4-shRNA” system. Genetics 193(1), pp. 51–61.

Sun, Z., Amourda, C., Shagirov, M., Hara, Y., Saunders, T.E. and Toyama, Y. 2017. Basolateral protrusion and apical contraction cooperatively drive Drosophila germ-band extension. Nature Cell Biology 19(4), pp. 375–383.

Takahashi, M., Takahashi, F., Ui-Tei, K., Kojima, T. and Saigo, K. 2005. Requirements of genetic interactions between Src42A, armadillo and shotgun, a gene encoding E-cadherin, for normal development in Drosophila. Development 132(11), pp. 2547–2559.

Takeichi, M. 2014. Dynamic contacts: rearranging adherens junctions to drive epithelial remodelling. Nature Reviews. Molecular Cell Biology 15(6), pp. 397–410.

Tamada, M., Farrell, D.L. and Zallen, J.A. 2012. Abl regulates planar polarized β-catenin tyrosine phosphorylation. Developmental Cell 22(2), pp. 309–319.

Tamada, M., Shi, J., Bourdot, K.S., et al. 2021. Toll receptors remodel epithelia by directing planar-polarized Src and PI3K activity. Developmental Cell.

Uechi, H. and Kuranaga, E. 2019. The tricellular junction protein sidekick regulates vertex dynamics to promote bicellular junction extension. Developmental Cell 50(3), p. 327–338.e5.

Vanderleest, T.E., Smits, C.M., Xie, Y., Jewett, C.E., Blankenship, J.T. and Loerke, D. 2018. Vertex sliding drives intercalation by radial coupling of adhesion and actomyosin networks during Drosophila germband extension. eLife 7.

Williams, M.L. and Solnica-Krezel, L. 2017. Regulation of gastrulation movements by emergent cell and tissue interactions. Current Opinion in Cell Biology 48, pp. 33–39.

Wodarz, A. 2008. Extraction and immunoblotting of proteins from embryos. Methods in Molecular Biology 420, pp. 335–345.

Wodarz, A., Ramrath, A., Kuchinke, U. and Knust, E. 1999. Bazooka provides an apical cue for Inscuteable localization in Drosophila neuroblasts. Nature 402(6761), pp. 544–547.

Yu, H.H. and Zallen, J.A. 2020. Abl and Canoe/Afadin mediate mechanotransduction at tricellular junctions. Science.

Zallen, J.A. and Wieschaus, E. 2004. Patterned gene expression directs bipolar planar polarity in Drosophila. Developmental Cell 6(3), pp. 343–355.

