## Supplemental Materials for "Src42A is required for E-cadherin dynamics at cell junctions during *Drosophila* axis elongation"

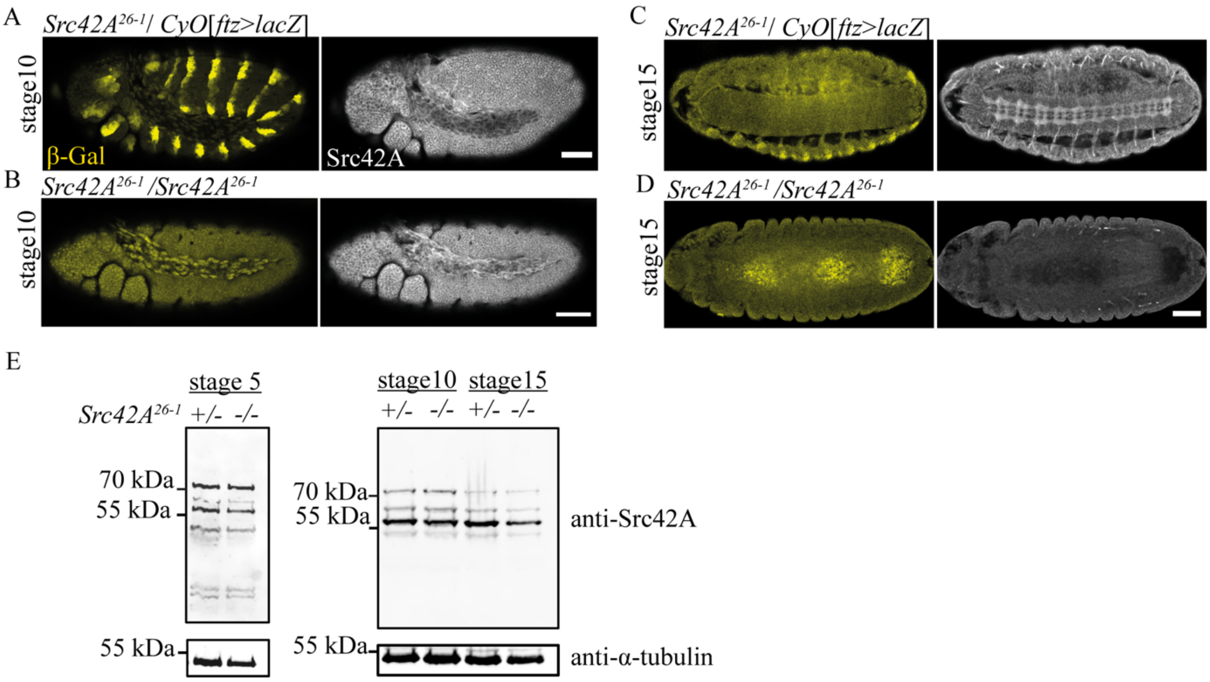


**Figure S1: Src42A antibody characterization in zygotic *Src42A$^{26\text{-}1}$* mutants**. **(A-D)** *Src42A$^{26\text{-}1}$*/ *CyO*[*ftz>lacZ*] is heterozygous viable mutant for *Src42A* used as a control and *Src42A$^{26\text{-}1}$*/ *Src42A$^{26\text{-}1}$* is homozygous zygotic mutant for Src42A; the *ftz>lacZ* balancer was used as a marker to discriminate the control and mutant embryos by anti-β-Gal staining. Scale bar represents 50µm. **(E)** SDS-PAGE followed by immunoblotting show maternal Src42A protein in zygotic mutants in different stages as marked above the blot, for each lane lysate of 20 embryos were loaded. +/- and -/- denotes heterozygous and homozygous mutants respectively. The predicted molecular weight for Src42A is 59kDa. An additional band observed approximately at 70kDa. The identity of the 70kDa band is unclear but note that the intensity of this band is reduced in later stage embryos mutant for *Src42A*. Anti-α-tubulin antibody was used as a loading control.

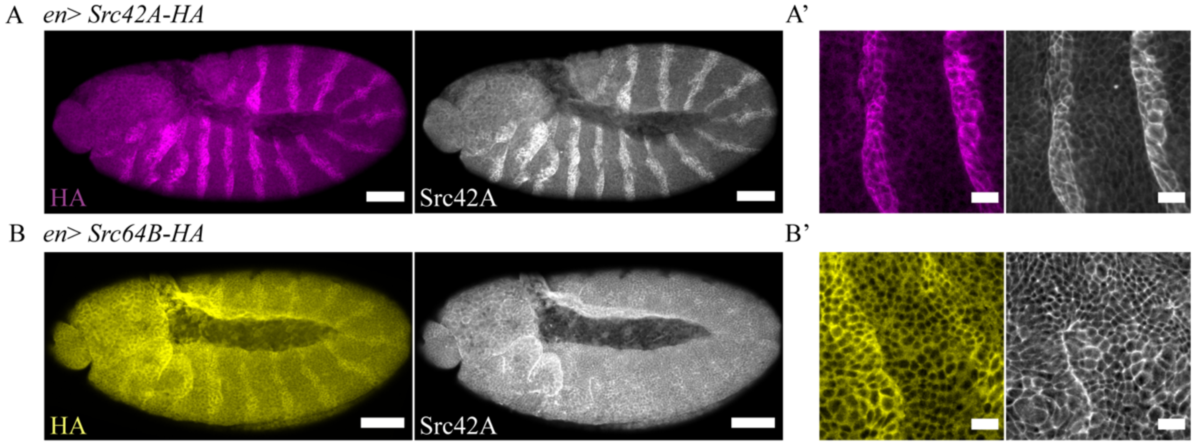


**Figure S2: Src42A antibody does not cross react with Src64B. (A)** Stage 10 embryos expressing *UAS::Src42A-HA* transgene using *engrailed>Gal4* driver flies. **(B)** Stage 10 embryos expressing *UAS::Src64B-HA*. Scale bars shows 50 µm. Images were taken at 25x magnification. **A'** and **B'** show 63x magnification images of *Src42A-HA* and *Src64B-HA* expression respectively. HA antibody staining marked in magenta and yellow and Src42A staining marked in grey. Scale bars corresponds to 10 µm.

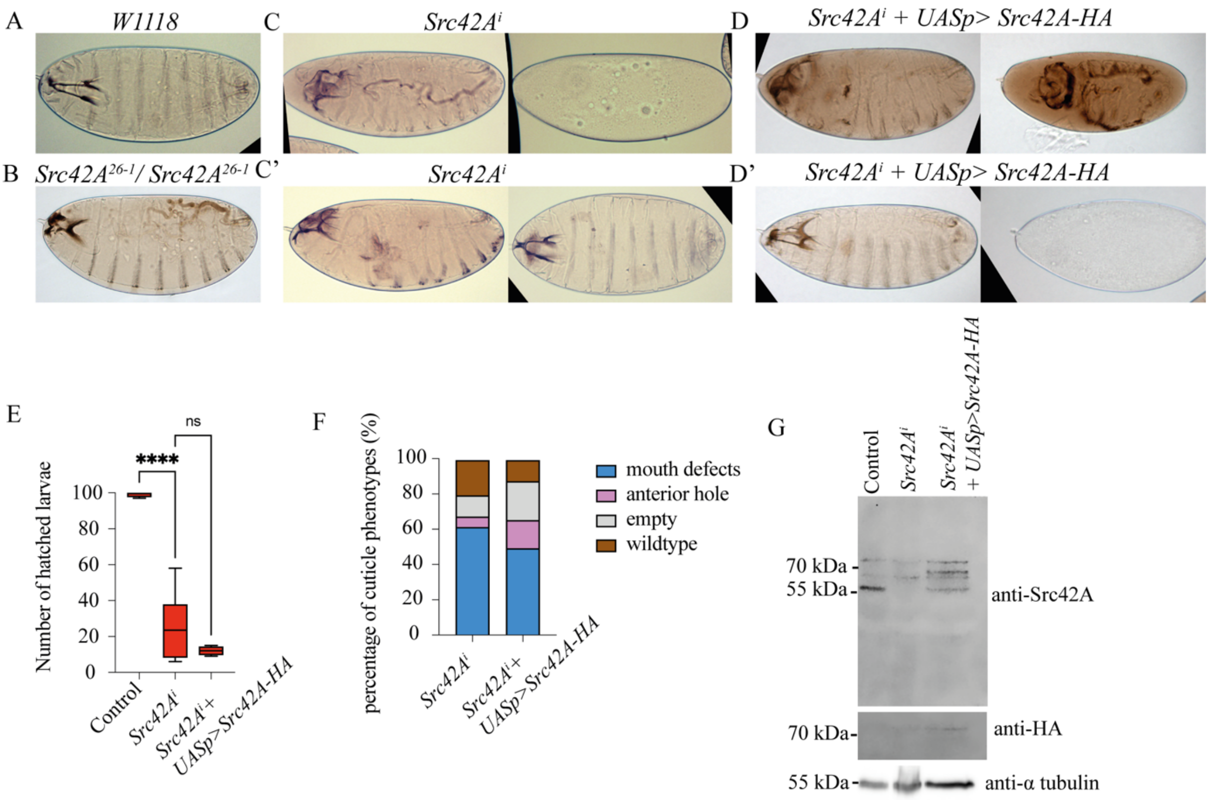


**Figure S3: *Src42A*$^{i}$ rescue attempt with *UASp>Src42A-HA*. (A, B)** Cuticle preparation from wild type (w$^{1118}$) and *Src42A* zygotic mutant (*Src42A*$^{26\text{-}1}$/*Src42A*$^{26\text{-}1}$) respectively. **(C, C')** *Src42A*$^{i}$ cuticles showing variable phenotypes. **(D, D')** Various cuticle phenotypes of *Src42A*$^{i}$*+UASp>Src42A-HA* embryos display similar effects as *Src42A*$^{i}$. **(E)** Hatching rate analysis of *Src42Ai* and rescue. Statistics were done using Dunnett's multiple comparison test shows significant difference among the mean values. The P value (0.1114) show no significant difference between *Src42A*$^{i}$ and *Src42A*$^{i}$*+UASp>Src42A-HA* rescue. **(F)** Cuticle classification from *Src42A*$^{i}$ and rescue analysis. **(G)** Western blot analysis showing *Src42A*$^{i}$ rescue analysis using *UASp>Src42A-HA*. Antibody against HA (haemagglutinin) shows very faint expression in higher molecular weight(~70kDa) range.
